# Population genetics in the early emergence of the Omicron SARS-CoV-2 variant in the provinces of South Africa

**DOI:** 10.1101/2023.02.09.527920

**Authors:** María Fernanda Contreras-González, Hugo G. Castelán-Sánchez, Erik Diaz-Valenzuela, Angélica Cibrián-Jaramillo

**Author notes:** María Fernanda Contreras-González, Hugo G. Castelán-Sánchez, Erik Díaz-Valenzuela, Angelica Cibrian-Jaramillo. Correspondence to: Angelica Cibrian-Jaramillo. These authors contributed equally to this word (María Fernanda Contreras-González, Hugo G. Castelán-Sánchez).

## Abstract

Population genetic analyses of viral genome populations provide insight into the emergence and evolution of new variants of SARS-CoV-2. In this study, we use a population genetic approach to examine the evolution of the Omicron variant of SARS-CoV-2 in four provinces of South Africa (Eastern Cape, Gauteng, KwaZulu-Natal, and Mpumalanga) during the first months before emergence and after early spread. Our results show that Omicron polymorphisms increase sharply from September to November. We found differences between SARS-CoV-2 populations from Gauteng and Kwazulu-Natal and viruses from the Eastern Cape, where allele frequencies were higher, suggesting that natural selection may have contributed to the increase in frequency or that this was the site of origin. We found that the frequency of variants N501Y, T478K, and D614G increased in the spike in November compared with other mutations, some of which are also present in other animal hosts. Gauteng province was the most isolated, and most genetic variation was found within populations. Our population genomic approach is useful for small-scale genomic surveillance and identification of novel allele-level variants that can help us understand how SARS-CoV-2 will continue to adapt to humans and other hosts.

## Introduction

Severe Acute Respiratory Syndrome Coronavirus 2 (SARS-CoV-2), which causes COVID-19, was first reported in Wuhan, China in December 2019. It has since spread to seven continents, and several variants have emerged during the two years of the pandemic. Variants Alpha (B.1.1.7), Beta (B.1.351), Gamma (P.1), Delta (B.1.617.2), and Omicron (B.1.1.519) have been reported as variants of concern (VOC) [1]. VOCs are associated with higher infectivity, disease severity, greater transmission efficiency in human populations [1], and a greater possibility of infecting other animal hosts [2].

Omicron is particularly interesting because it emerged from a sudden COVID-19 wave in South Africa and quickly spread worldwide [4]. It includes the B.1.1.529 lineage and lineages descended from it, as defined by WHO [1, 4, 5]. The Omicron spike mutation is believed to favor better transmission to humans, and it is increasingly clear how it arises among many hosts within the population and whether they evolve within a few individuals with intrahost variants [7, 8, 9]. Furthermore, fixed mutations in the spike region are thought to be adaptive, yet they vary spatially and temporally between geographic regions [11, 12], which has been used to monitor the transmission and spread of the virus in different populations [13].

Population genetics can provide early information on the emergence and maintenance of alleles that may become key mutations for virus adaptation to new hosts or conditions. Even before they form a distinct phylogenetic lineage or clade, or their status as variants of concern is determined, genetic signatures at the population level can indicate potentially novel variants of concern [3]. The possibility that Omicron has a different animal origin is also intriguing, and could be explored using a population genetics approach.

Population genetic approaches have been used to monitor the frequency of mutations, alternative alleles, lineage spread and population structure, and evolutionary dynamics of SARS-CoV-2 [14, 15]. These approaches have allowed for the detection of recurrent mutations and facilitated the visualization of genomic variation and lineage spread [14]. Previous population genetic studies of polymorphisms have revealed that SARS-CoV-2 genomes exhibit regional population structure in SARS-CoV-2 populations [13]. Our previous population-level study was able to identify variants associated with asymptomatic patients that were not detected in phylogenetic studies [3].

In this study, we measure allele frequencies of genomes sequenced from patients in South Africa at the onset of Omicron to understand its origins and describe its early evolution in four provinces of South Africa: Eastern Cape, Gauteng, KwaZulu-Natal, and Mpumalanga, using population genomics. Our results show that Omicron polymorphisms increased over time (September to November). These polymorphisms were mainly concentrated in the spike protein, but varied between provinces and months. We found that the distribution of alternative alleles to the canonical SARS-CoV-2 from Wuhan and the alpha variant reported as a possible origin [6] was higher in September before the appearance of Omicron. In contrast, the frequency of the alternative alleles decreased after the appearance of Omicron in November.

We also observed differentiation between SARS-CoV-2 populations from Gauteng and KwaZulu-Natal compared to viruses from the Eastern Cape, where allele frequencies were higher, suggesting that the increase in frequency was due to natural selection. We found that the Eastern Cape had a low level of low polymorphisms potentially influenced by balancing selection. There is high genetic variation within populations (95.8%), with Gauteng as the most isolated population. Our results describe how polymorphisms spread and give rise to new variants in SARS-CoV-2.

Population genetics can provide useful information about the evolution of Omicron and other virus variants, as well as inferences about the genetic diversity of the virus in space and time. This information is crucial for understanding the emergence of new variants, predicting their impact on human health and controlling the spread of the virus.

## Materials and Methods

### Variant Calling

A catalog of 504 paired-end sequencing libraries collected from human hosts between September and November 2021 was retrieved from the Sequence Read Archive (SRA) of the National Center for Biotechnology Information (NCBI) on December 6, 2021. These libraries were obtained using real-time polymerase chain reaction (RT-PCR) and were originally isolated from oropharyngeal swabs from 504 patients of indeterminate sex.

We followed a population-genetic strategy previously used for SARS-CoV-2 population-level descriptors [3]. First, the raw reads from these 504 libraries were first processed using the fastp software version 0.20 with default parameters for filtering by quality and length and removing adapters [16]. The clean reads were then mapped to the SARS-CoV-2 reference genome (NC_045512.2) using BWA v0.7.15 [17]. The SAM alignments were then sorted and converted to BAM files using samtools [18]. Subsequently, the BAM alignments were used for genetic variant detection using freebayes v.1.2 [19]. The individual variant call format files (VCFs) were then merged into a single file. Variants were then filtered and parsed using the R package tidyverse 1.3.1 to meet the following criteria: a minimum quality score of 30, a minimum of 20 reads representing each locus, only one alternative version in the host individual (biallelic), and a frequency of at least 0.05 for each variant in all samples. Of the 504 libraries, 448 passed the filters, with samples from the four provinces of South Africa distributed as follows: Eastern Cape (85 individuals), Gauteng (142 individuals), KwaZulu-Natal (216 individuals), and Mpumalanga (5 individuals).

### Polymorphism Sites Frequency

To determine the frequency of SARS-COV −2 mutations in the South African population in September and October 2021, nucleotide variants and indels were assessed using a strategy that mapped raw data to the reference genome (NC_045512.2). To examine the intrahost variation of SARS-CoV-2 reads, the frequency of alternative alleles was calculated [3]. i.e., the proportion of total reads that are derived alleles-either nucleotide variants or indels-in each library. This measure was calculated using R functions developed inhouse. i.e., the proportion of total reads that are derived alleles-either nucleotide variants or indels-in each library.

To assess how genetic variation changed over time, the dataset was split by month: September (110 individuals), October (166 individuals), and November (172 individuals). And to analyze genomic changes per province, the dataset was split by province: Eastern Cape (85 individuals), Gauteng (142 individuals), KwaZulu-Natal (216 individuals), and Mpumalanga (5 individuals). Mpumalanga was excluded due to small sample size. Allele frequencies per dataset were recalculated to fit the sample size of each dataset. The polymorphism frequencies and the mean allele proportions were plotted as a lollipop plot from the R package ggplot2 3.3.5. The correlation between the read depth and the discovery of nucleotide polymorphisms was also tested using cor.test function, with Pearson method, from the stats R package.

The polymorphisms found were compared against the defining mutations of some VOCs as they were reported in the Cov-Lineages webpage (https://cov-lineages.org/constellations.html) on April 13, 2022 [20]. The main VOC, in which its mutations were contrasted, was Omicron lineage prior to splitting into sublineages [4]. Other VOCs defining mutations were compared, due to their similarities with Omicron [5], these were: Alpha [21] and Delta [5] variants. The polymorphisms site frequency was recalculated for each subset of data considering their respective sample size: either by province (Eastern Cape: 85, Gauteng: 142, and KwaZulu-Natal: 216) or by month (September: 110, October: 166, and November: 172). A Kolmogorov-Smirnov test from R package stats 4.2.0 was used to perform pairwise comparisons of the distributions of each dataset grouped by month and province. This information was displayed as a violin plot and box plot using ggplot 3.3.5.

### Genetic structure

Single nucleotide polymorphisms (SNPs) were obtained from 504 libraries (excluding samples from Mpumalanga) and converted to allelic dosage. This allelic dosage consisted of a matrix of individuals and alternative alleles, containing the number of alternative alleles in each individual [22]. Given the haploid nature of SARS-CoV-2, the genotypes were transformed into counts as follows: 1/1 and 0/1, meant presence and were interpreted as 1; NA meant that the SNP was not found, thus was 0 [22]. Allele dosages were separated by province (Eastern Cape, Gauteng and KwaZulu-Natal). The separated doses were used to calculate nucleotide diversity (pi), Tajima’s D as a proxy for selection pressure, and the F statistic to assess the extent of differentiation and correlation of randomly selected alleles within the same subpopulation relative to the overall population [23], these parameters were calculated with the R package hierfstat 0.5-10. With this R package and using the same steps, each position that had an SNP was explored in order to calculate their respective Fst value.

Population structure assessment with an AMOVA of 504 libraries was performed using the R package poppr 2.9.3 and adegenet 2.1.5. The genind object relies mainly on a matrix of individuals and all alleles of a locus containing their counts, corresponding to ploidy [24]. Thus, the reference alleles were given a value of 0 if they were homozygous for the alternative allele “1/1”; in the other cases, the reference alleles had a value of one. The same was true for the alternative alleles, which had a value of zero if there was no data on them “NA”, the remaining cases “1/1” or “0/1” were interpreted as one. The structure given by the province AMOVA was then calculated using the Genind object by province [24]. A Monte Carlo test was performed to test for significance [24].

## Results

### Identification of genetic variants within the South African SARS-CoV-2 genome

We identified positions along the SARS-CoV-2 genome in samples from the South African provinces that showed divergence compared with the reference genome. Before frequency filtering, there were a total of 2376 nucleotide variations in the 503-library dataset. After filtering, 448 libraries remained with 91 nucleotide polymorphisms. Of the 91 polymorphisms identified, 28 belonged to the Omicron/B.1.1.529 lineage, 13 to the delta and 2 to the alpha constellation of defining mutations (**Figure 1**). Only one mutation, T478K on the spike protein, was part of both the delta and omicron mutational constellations (**Figure 1**). The two alpha-defining mutations, N501Y and P681H, occurred in the entire Omicron lineage (**Figure 1**). The Omicron mutations considered are those shared by all Omicron sublineages (2022).

**Figure 1.**
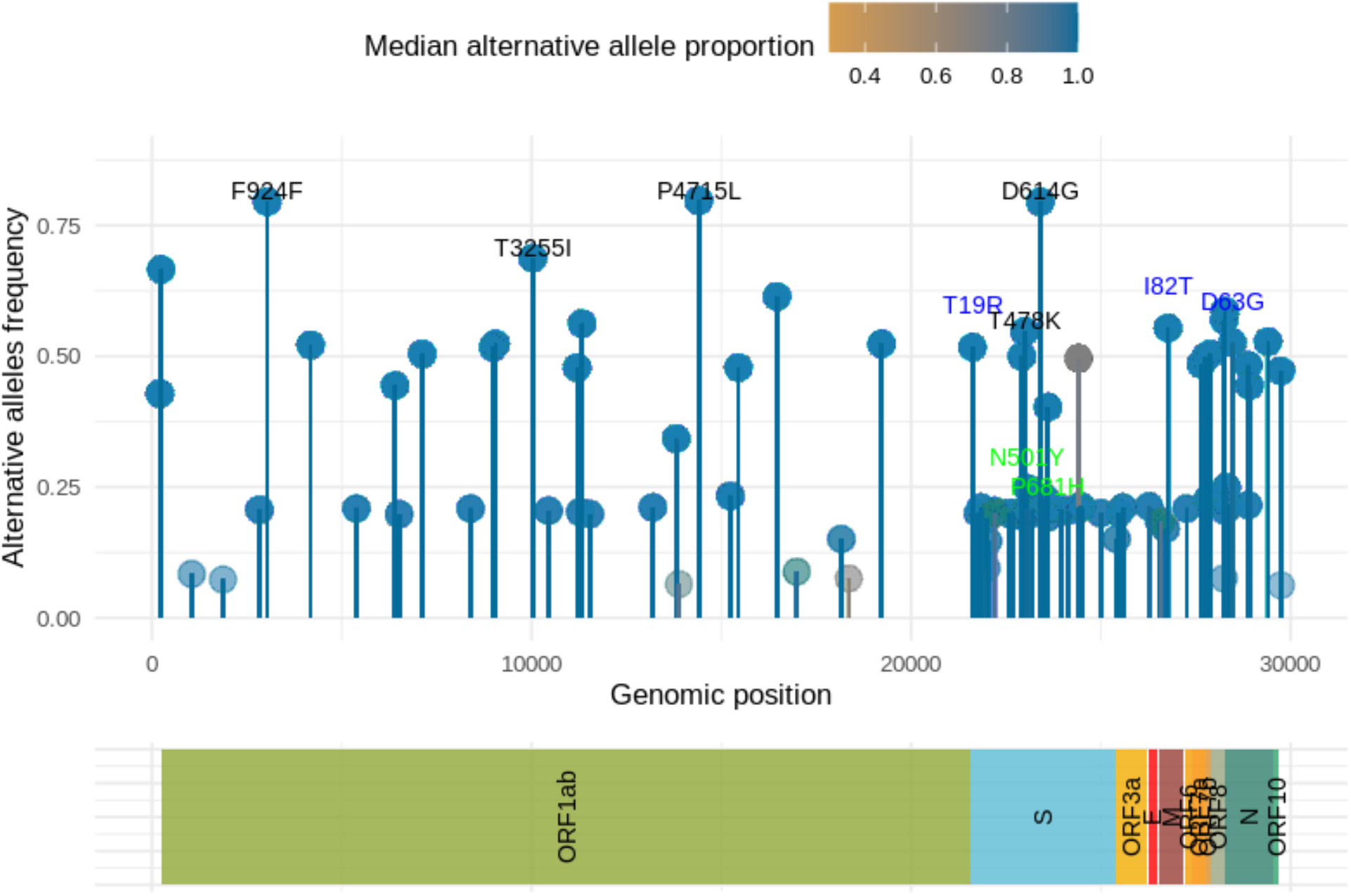
Genomic changes in SARS-CoV-2 reads in South Africa in 2021. Nucleotide variants and indels found in SARS-CoV-2 sequences. The y-axis shows the frequency of each change, the x-axis shows the genomic position, and the color code indicates the median proportion of alternative alleles in the entire data set. The words above each lollipop indicate the mutation of a constellation variant with a frequency of ≥0.5 (except for Alpha mutations) in the samples, Omicron in black, Delta in blue, and Alpha in green, although not all are shown due to overlap. The mutations names are displayed as their predicted amino acid changes.

Subsequently, the South African dataset was divided into months (September, October, and November), and an increase in common mutations of the Omicron lineage was observed in November (**Figure 2**). The following number of B.1.1.529 lineage mutations were detected per month in the 448 libraries from the four provinces: eight in September, eight in October, and 28 in November (**Table 1**). This is a remarkable increase, as it corresponds to a substitution rate of 2.675 x 10-03 per site per year.

**Figure 2.**
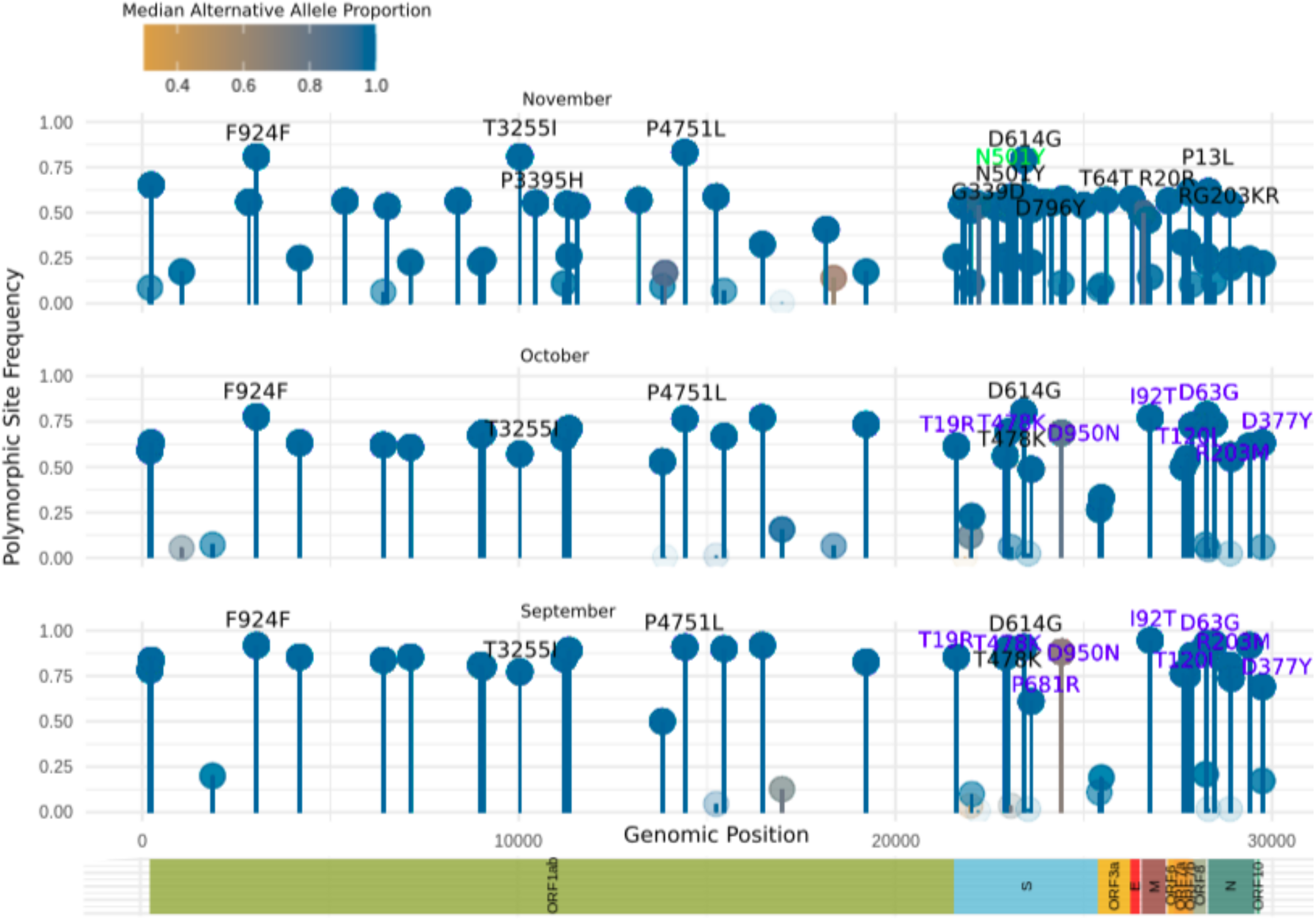
Mutations in SARS-CoV-2 reads by month in four South African provinces. Each graph shows the frequency of each change on the y-axis, the genomic position on the x-axis, and the color code indicates the median proportion of alternative alleles in the entire data set. The words above each lollipop indicate the mutation of a constellation variant with a frequency of ≥ 0.5 in the samples (except for Alpha mutations), in black Omicron and in blue Delta, not all are shown due to overlap. The months shown are: A) September, B) October, and C) November. The mutations names are displayed as their predicted amino acid changes.

**Table 1.**
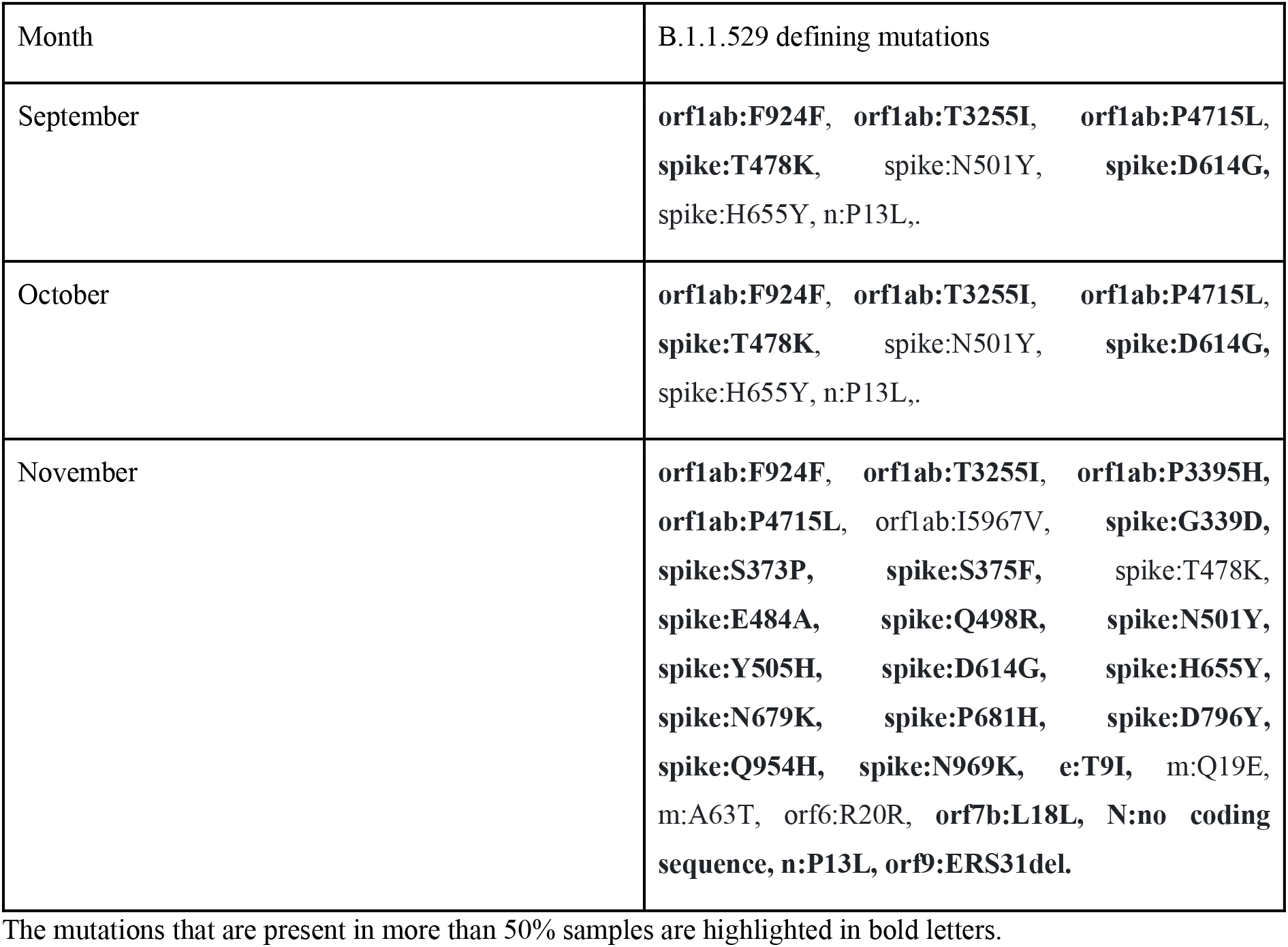
Defining mutations per month of the Omicron B.1.1.529 lineage constellation present in the SARS-CoV-2 genomes in the provinces of South Africa 2021.

The frequency of Omicron lineage mutations changed over time (**Figure 2**). Omicron mutations in > 50% of samples increased over the months, from five in September and October to 28 in November. Delta mutations present in > 50% of samples were dramatically reduced (**Figure 2**), from 11 mutations in September to nine in October and none in November. Mutations with reduced frequency in samples included the T478K mutation in spike, which is also present in Omicron (**Supp. Table 1**). Two alpha-defining mutations were present, originating from the spike region and shared with Omicron. The N501Y mutation is present in all three months and the P681H mutation is present only in November; both increase their occurrence in November to more than 50% of the samples (**Supp. Table 2**).

In total, Omicron B.1,1,529 mutations arose rapidly, as shown by our estimates of substitutions per site. Genome-wide substitutions per site per year were 2.6×10^-3^ (28-8 substitutions /29903 bases / 0.25 years = 20/29903/0.25 = 0.002675 = 2.6×10^-3^), which is higher than the previously estimated rate of substitution per site per year of 8.90e-4 [25]. In the spike region, there were 1.15×10^-2^ substitutions per site per year (15^-4^ substitutions / 3822 sites in the spike region /3 months = 11 / 3822 / 0.25 = 0.0115 = 1.15×10^-2^), which is higher than the range of 1.06 × 10^-3^ and 1.69 × 10^-3^ previously reported for this region for sequences from different regions worldwide [26].

Considering the mutations of the SARS-CoV-2 variant by province, mutations common to the B.1.1.529 lineage are found in all provinces (**Table 2, Figure 3**), although the number of mutations varies from province to province (Figure 3). There were 28 mutations in Gauteng, 28 in KwaZulu-Natal, and eight in Mpumalanga. In addition, four mutations of Omicron are present in more than 50% of the samples in each province: orf1ab:F924F, orf1ab: T3255I, orf1ab: P4715L, and spike: D614G.

**Figure 3.**
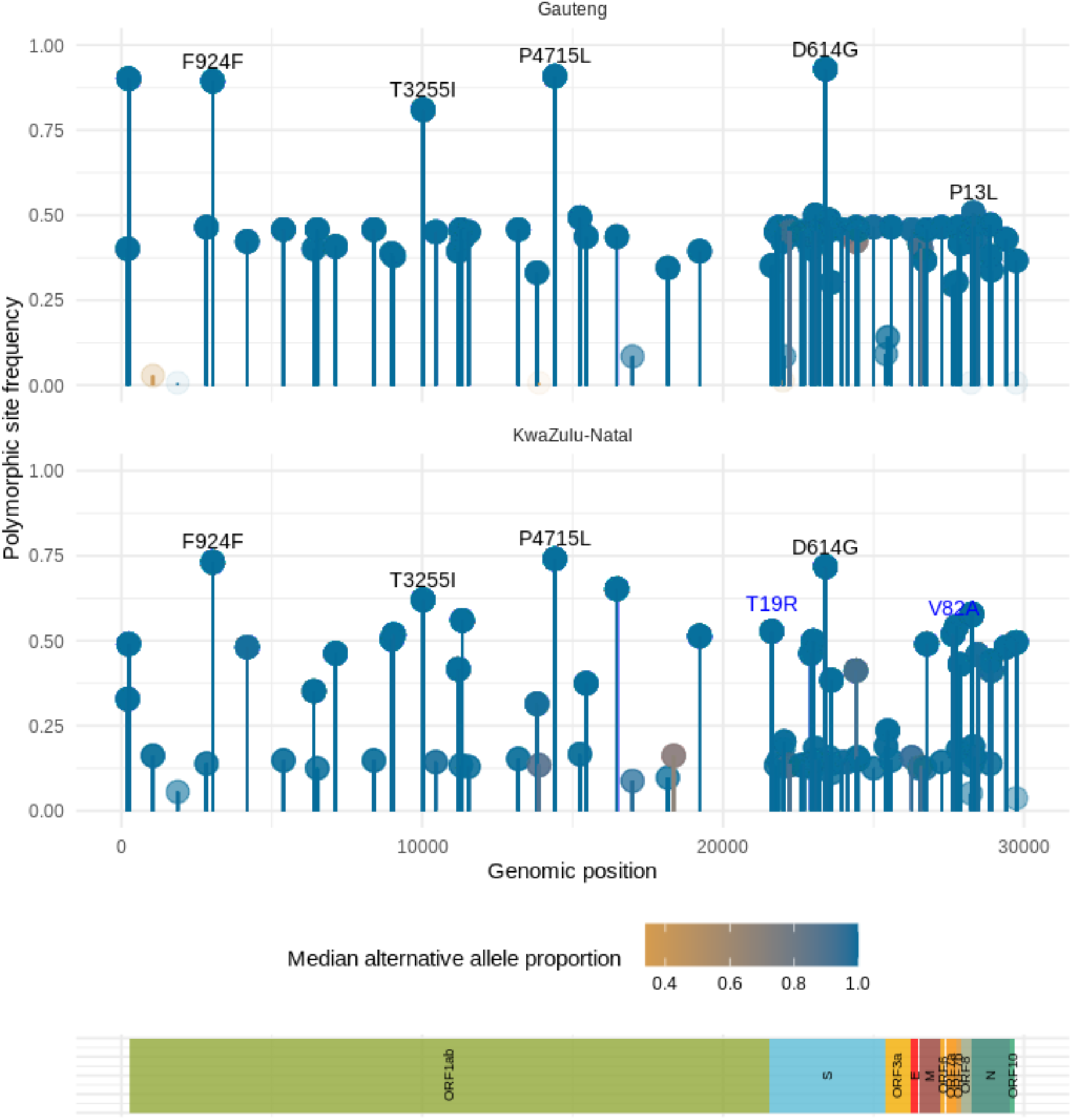
Mutations in SARS-CoV-2 reads from South Africa during September-November 2022 in two provinces. Each plot shows the frequency of each change on the y-axis, the genomic position on the x-axis, and the color code indicates the median proportion of alternative alleles in the entire data set. The words above each lollipop indicate the mutation of a constellation variant with a frequency of ≥ 0.5 in the samples, in black Omicron and in blue Delta, not all are shown due to overlap. The provinces shown are: A) Gauteng and B) KwaZuluNatal. The mutations names are displayed as their predicted amino acid changes.

**Table 2.** Defining mutations per month of the Omicron B.1.1.529 lineage constellation present in the SARS-CoV-2 genomes in the provinces of South Africa 2021.

| Month | B.1.1.529 defining mutations |
| --- | --- |
| September | **orf1ab:F924F**, **orf1ab:T3255I**, **orf1ab:P4715L**,<br>**spike:T478K**, spike:N501Y, **spike:D614G**,<br>spike:H655Y, n:P13L. |
| October | **orf1ab:F924F**, **orf1ab:T3255I**, **orf1ab:P4715L**,<br>**spike:T478K**, spike:N501Y, **spike:D614G**,<br>spike:H655Y, n:P13L,. |
| November | **orf1ab:F924F**, **orf1ab:T3255I**, **orf1ab:P3395H**,<br>**orf1ab:P4715L**, orf1ab:I5967V, **spike:G339D**,<br>**spike:S373P**, **spike:S375F**, spike:T478K,<br>**spike:E484A**, **spike:Q498R**, **spike:N501Y**,<br>**spike:Y505H**, **spike:D614G**, **spike:H655Y**,<br>**spike:N679K**, **spike:P681H**, **spike:D796Y**,<br>**spike:Q954H**, **spike:N969K**, **e:T9I**, m:Q19E,<br>m:A63T, orf6:R20R, **orf7b:L18L**, **N:no coding**<br>**sequence**, n:P13L, **orf9:ERS31del**. |

The mutations that are present in more than 50% samples are highlighted in bold letters.

### Allele frequency distribution shows higher diversity within-population than among populations for most sites

During the spread of the Omicron lineage in South Africa, a number of predominant mutations of the SARS-CoV-2 lineage changed according to province and time. We examined the distribution of the genome-wide proportion of alternative alleles of SARS-CoV-2 libraries within and between provinces and their changes during the three-month period. The proportion of alternative alleles was higher in September than in November, when the distribution of alternative alleles was narrower.

Comparing this distribution by province (**Figure 4**), Kwazulu-Natal showed a predominantly left-skewed distribution, indicating a high density of low and intermediate frequency polymorphisms in this province. The distribution of polymorphisms in Gauteng more closely resembles a Gaussian distribution with density in the middle. The Eastern Cape has a bimodal distribution indicating a high density of high frequency polymorphisms (Figure 4). A Kolmogorov-Smirnov test showed that all the distributions of the provinces were statistically significantly different from each other, with a p-value of < 2.2e-16. The main difference arose from the distances, with the largest most significant difference between KwaZulu-Natal and Eastern Cape (D=0.88631), followed by Eastern Cape and Gauteng (D=0.81217); finally, the smallest differences were between KwaZulu-Natal and Gauteng (D=0.3215).

**Figure 4.**
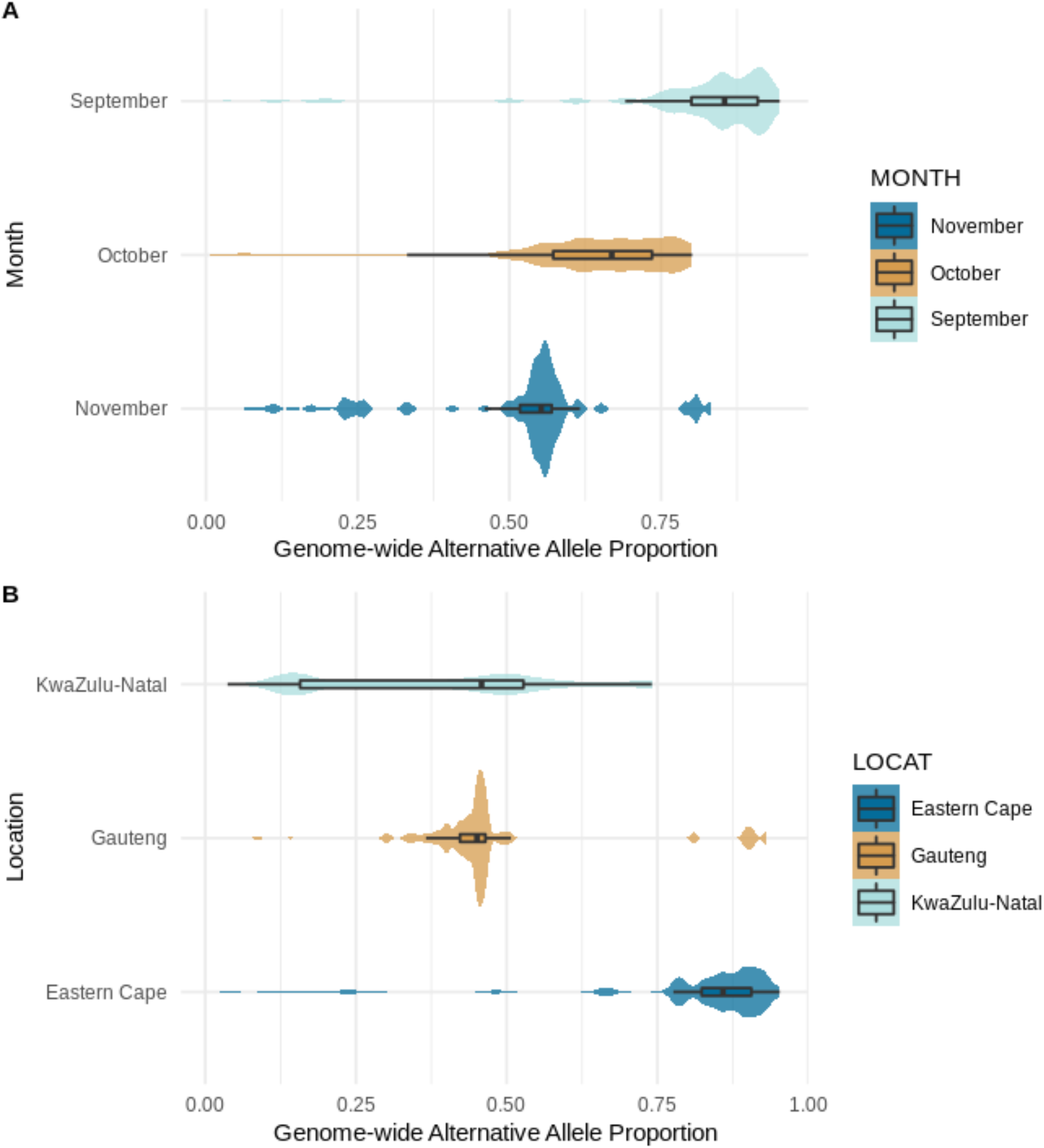
Genome-wide allele proportion grouped by month and location. Site frequency spectra are shown for all months (September, October, and November) and for three provinces (KwaZulu-Natal, Gauteng, and Eastern Cape) to compare genome-wide nucleotide changes.

The increased allelic frequency of some mutations and the skewed frequencies could be due to increased fitness of some virus populations by natural selection. Therefore, we tested for the presence of natural selection using the Tajima test. The results of the Tajima test show the presence of positive selection for the three provinces, suggesting that balanced selection favors the maintenance of multiple alleles in these populations, at higher frequencies than would be expected based on genetic drift alone (**Table 3**).

Regarding the average nucleotide diversity (pi) for the whole genome length (29903 nucleotides, accession number: NC 045512), Gauteng had the highest value (pi = 0.00083414) and Eastern Cape the lowest value (pi = 0.00053164), suggesting that the virus might have different effective population sizes in different regions. According to the pairwise FST estimates of population differentiation, all three sites had some degree of differentiation, with Gauteng having the lowest gene flow among the other sites (**Table 3**).

**Table 3.** Genetic diversity measures.

| Province | Pi from dosage | Tajima's D |
| --- | --- | --- |
| **Eastern Cape** | 0.0005383482 | 3.68537 |
| **Gauteng** | 0.000802908 | 2.872843 |
| **KwaZulu-Natal** | 0.000586203 | 1.48053 |

**Pairwise *F*sts**
|  | **Eastern Cape** | **Gauteng** | **KwaZulu-Natal** |
| --- | --- | --- | --- |
| **Eastern Cape** | - | 0.13726290 | 0.06767264 |
| **Gauteng** | 0.13726290 | - | 0.04674843 |
| **KwaZulu-Natal** | 0.06767264 | 0.04674843 | - |

**Global *F*sts**
| **All** | **Eastern Cape** | **Gauteng** | **KwaZulu-Natal** |
| --- | --- | --- | --- |
| 0.08607307 | 0.2359730 | -0.1428762 | 0.1651225 |

However, the global FST values show a different degree of correlation of randomly selected alleles within the same subpopulation compared to the overall population, with Gauteng and KwaZulu-Natal showing more similar allele frequencies within populations than the Eastern Cape in comparison. This is consistent with what we observed for the overall distribution of allele frequencies in the frequency spectra of polymorphic sites overall and per month (**Figure 4, Supp. Figure 4**).

The FST value was calculated for each genomic position that had a SNP (**Figure 5**). Global FST values ranged from −0.002 to 0.152, while provincial FST values had a wider range: Eastern Cape from −1.201 to 1, Gauteng from −1.019 to 1, and KwaZulu-Natal from −1.763 to 0.534. Of the twelve highest FST values, seven belonged to mutations defining the entire Omicron lineage (**Figure 5, Supp. Table 5**): N:no-cds or nuc:A28271, spike:N679K, orf6:R20R or nuc:A27259C, spike:Q954H, spike:G339D, spike:S375F, spike:D796Y. Among them there are also two shared only by BA.1 and BA.2 Omicron sublines. It is worth noting that the region with the highest number of Omicron defining mutations with a non-zero FST value is spike.

**Figure 5.**
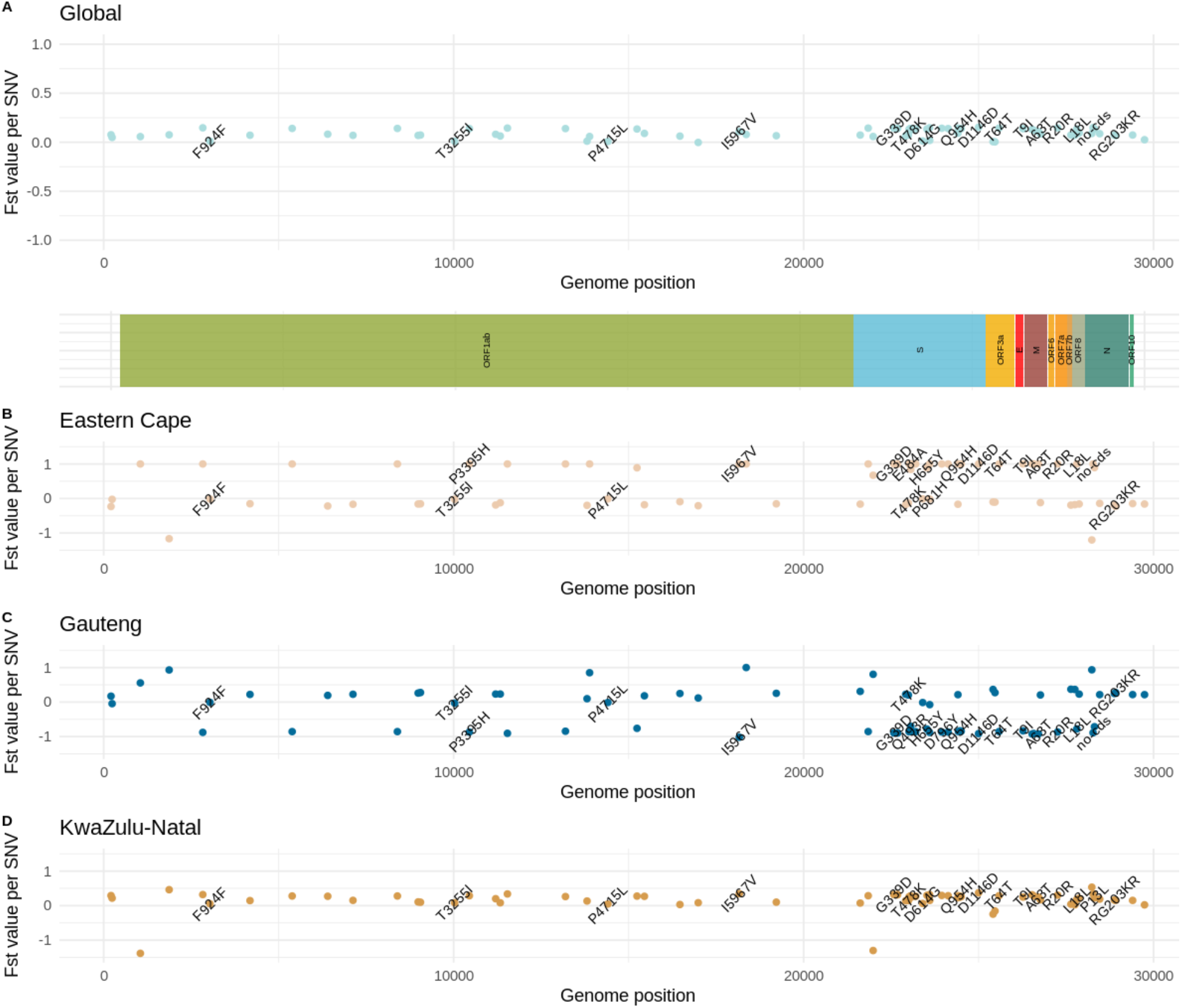
FST values per SNV found. A) The Fst value was calculated for each SNV of all the libraries of South Africa between September and November 2021. Fst values calculated for each SNV for each province: B) Eastern Cape, B) Gauteng and C) KwaZulu-Natal. Omicron lineage defining mutations that have FST values different to zero were added as black labels.

According to AMOVA, the frequency of alternative alleles varied significantly more between samples than between provinces and between September and November (**Table 4**). The greatest variation was found in the provinces, and the frequency of the alternative alleles was significantly different in at least one of these communities. According to the AMOVA results, differences between specific individuals within populations (95.8% of total variation) were more significant than differences between different provinces (4.2% of total variation). In addition, the population differentiation (phi) statistic from a Monte Carlo test showed considerable variation between populations, as it did not match the predicted distribution (**Supp 5. Figure, Table 4**).

**Table 4.** Monte Carlo test between provinces.

| **Monte Carlo test between samples** |  |  |  |
| --- | --- | --- | --- |
| **Observation** | **Std. Obs.** | **Alter** | **P-value** |
| 1.196231 | 8.65880203 | greater | 0.01 |

## DISCUSSION

We used a population genomics approach to analyze the Omicron variant as it emerged in South Africa.We focused on the provinces of KwaZulu-Natal, Gauteng, Eastern Cape, and Mpumalanga, the only South African provinces where paired SARS-CoV-2 libraries were present in the months following the discovery of Omicron. We monitored genome-wide changes in allele frequency in SARS-CoV-2 reads from South Africa within three months comprising the first report of Omicron VOC [27, 31].

The frequency of mutations defining the Omicron lineage increased from September to November, resulting in VOC Omicron. The most frequent mutations of known VOCs, such as Spike:T478K, occurred in Delta and Omicron from September to November 2021. Gauteng was the province where Omicron lineage-defining mutations occurred more frequently, consistent with the fact that Gauteng was the first province where multiple Omicron-like sequences were discovered [4].

The emergence and establishment of overall Omicron mutations in the four provinces of South Africa between September and November 2021 were variable and appeared to be sitespecific. However, they all had in common that they occurred rapidly, as indicated by their allele frequencies and substitution rates. In the full dataset, there is an increase of 20 mutations within three months, rather than the expected 6.5 mutations calculated for three months using the previously genome-wide estimated substitution rate of 8.90e-4 per site per year [25]. It is worth noting that the substitution rate of 8.90e-4 per site per year was estimated for a single strain [25], which may indicate that more than one strain was active in the four provinces during this period.

There were eight defining mutations shared by the entire Omicron lineage and present in all provinces and all months: orf1ab:F924F, orf1ab:T3255I, orf1ab:P4715L, spike:T478K, spike:N501Y, spike:D614G, spike:H655Y, and n:P13L. These eight common mutations of the Omicron lineage shared some common features: Most of them are nonsynonymous mutations, with the exception of orf1ab:F924F or nuc:C3037T; and most are in either orf1ab or spike. Four of these eight mutations are part of the mu constellation of defining mutations, namely: orf1ab:F924F, orf1ab:T3255I, spike:N501Y, and spike:D614G [20]. Two of them are unique to Omicron, namely orf1ab:P4751L and spike:H655Y. For some of these four common mutations in the Omicron lineage, there is evidence linking them to increased fitness, such as spike:D614G by enhancing transmission [28], epitope loss that could lead to immunological scape, such as orf1ab:P4715L [29]; reduction in severity of COVID-19 cases, such as orf1ab:T3255I [30]. Although the synonymous mutations orf1ab:F924F or nuc:C3037T do not cause changes in any protein, there is a strong allelic association with other non-synonymous mutations such as orf1ab:P47I15L and spike:D614G [31, 32], which may contribute to the maintenance of virus integrity.

Two of these common Omicron-defining mutations, spike:N501Y and spike:H655Y, were found to increase SARS-CoV-2 infectivity in mice [7, 8, 9], which may partially explain the origin of some of these mutations that have no precedents in previous VOCs, such as H655Y [20]. Overall, these molecular spectrum analyses have shown that it is unlikely that Omicron mutations were acquired in humans before the outbreak, and our results may confirm this idea for these two changes that have been present since September.

Most variation (89.8%) occurred within the populations studied, although at least one province had significantly different proportions of alternative alleles. This supports the hypothesis that some individuals have high diversity of SARS-CoV-2 genomes and that multiple viral subpopulations evolve simultaneously in the areas studied. Our nucleotide diversity (pi) values also show high diversity of SARS-CoV-2 sequences due to balancing selection [35]. This high diversity of SARS-CoV-2 genomes due to the high number of mutations can be compensated by recombination to avoid deleterious mutations [37] or by template switching in the positive-stranded RNA virus SARS-CoV-2 [34o].

Indicators of genetic structure among populations showed differences, suggesting high genetic variation within populations. Genetic differentiation values other than zero could indicate selection pressure in viral populations [36], which is consistent with Tajima’s D results and FST values. The region with more SNVs with high FST values was spike, and the omicron-defining mutation. This suggests that there is specific selection pressure that may have driven Omicron selection; other regions that also had some high FST values were Nuc and Orf6, which may also be relevant contributors. It is also worth noticing that most of the Omicron mutations that had values different to zero are still up-to-date (February 2023) part of the new Omicron sublineages, which again support the role of natural selection [20].

The genome-wide fine-scale distribution of nucleotide variants, the frequency spectra for three provinces (Eastern Cape, Gauteng, and KwaZulu-Natal) differ. Eastern Cape exhibits skewness toward the most common nucleotide variants and is the one with the greatest distance from the other provinces. This could be explained by the lack of data in November, or because the Eastern Cape did not have as many Omicron cases as Gauteng or KwaZulu-Natal [4]. As time progresses, less frequent nucleotide variants in September shift to the most frequent nucleotide variants at the end of the year. A positive Tajima’s D suggests that this increase in allele frequencies is indeed due to balanced selection [33, 34, 35].

In conclusion, population genetic analysis allowed us to observe how mutation frequencies changed over time. Our results show that omicron polymorphisms increased over time (September to November), and SARS-CoV-2 populations from Gauteng and Kwazulu-Natal differed markedly in allele frequencies from viruses from the Eastern Cape. The evolution and development of new SARS-CoV-2 variants could be understood by population genetic studies of viral genomes. Future population genomic analyses are important for genomic surveillance and management of potentially new viral strains. Analysis of genetic variants of viruses using a population genetics approach is important for tracking the minor mutation spectrum in RNA viruses.

## Acknowledgments

This work was partially funded by Conacyt, grant No. 313075 to ACJ.

## Supplementary Information

The analyses and the information used to do it can be found in this GitHub repository: https://github.com/fglez/omigenpop.

Supplementary figures and table can be downloaded from here: https://drive.google.com/file/d/16CUjUksvvVlQsOCIgY1ng7Iqzm7XgOQv/view?usp=sharing

